# A Goldilocks principle of JAK/STAT signaling governs airway epithelial homeostasis and stress adaptation in *Drosophila*

**DOI:** 10.1101/2025.05.20.655050

**Authors:** Xiao Niu, Christine Fink, Kimberley Kallsen, Leizhi Shi, Viktoria Mincheva, Sören Franzenburg, Ruben Prange, Jingjing He, Anita Bhandari, Iris Bruchhaus, Holger Heine, Judith Bossen, Thomas Roeder

## Abstract

Airway epithelia must maintain barrier integrity while continuously adapting to changing environmental conditions. This requires key signaling pathways in airway epithelia to operate within a tightly controlled functional range. Here, we show that airway epithelial homeostasis in *Drosophila* depends on balanced JAK/STAT signaling. Basal pathway activity is constitutively present in differentiated epithelial cells and is necessary for cell survival, epithelial integrity and stress resistance. In accordance with this role as a stress responsive regulator in epithelial biology, environmental stressors, including hypoxia, cigarette smoke, and cold exposure, induce JAK/STAT signaling. In contrast, sustained pathway activation is associated with airway remodeling characterized by epithelial thickening, luminal narrowing, and altered cellular organization. Transcriptomic analysis reveals that sustained activation is associated with a coordinated epithelial stress program integrating immune signaling, proteostasis, and metabolic adaptation. Cross-species comparisons with murine and human datasets suggest that key aspects of this response are conserved. Pharmacological inhibition demonstrates that remodeling depends on continued pathway activity and can be partially reversed *in vivo*. Together, our findings support a model in which airway epithelial homeostasis depends on maintaining JAK/STAT signaling within a defined functional range. This “Goldilocks” principle provides a conceptual framework for understanding how epithelial stress responses are balanced under physiological and pathological conditions.

**Graphical abstract:** 

## Introduction

Airway epithelia form the first barrier between the organism and the external environment and are therefore continuously exposed to environmental stressors, including pathogens, pollutants, and fluctuations in oxygen availability[1, 2]. Maintaining epithelial integrity under these conditions requires the coordinated regulation of signaling pathways that govern epithelial survival, stress responses, and tissue architecture[3, 4]. Dysregulation of epithelial signaling can lead to pathological remodeling processes that contribute to chronic airway diseases, including asthma and chronic obstructive pulmonary disease [5-7]. Understanding how epithelial signaling pathways maintain airway homeostasis while enabling adaptive responses to environmental stress, therefore, represents a fundamental challenge in lung research.

Among the signaling pathways implicated in epithelial regulation, the JAK/STAT signaling pathway plays a central role in epithelial stress responses and tissue remodeling across metazoans[8, 9]. In mammalian airways, JAK/STAT signaling is activated by inflammatory cytokines and has been implicated in epithelial proliferation, immune responses, and airway remodeling during chronic lung disease[9-11]. However, despite extensive work in mammalian systems, it remains unclear how JAK/STAT signaling contributes to airway epithelial homeostasis under physiological conditions and how different levels of pathway activity affect epithelial structure and function[12-15][4].

The *Drosophila* tracheal system provides a powerful in vivo model to study airway epithelial biology. The fly airway epithelium shares key structural and functional features with mammalian airway epithelia while offering unparalleled genetic accessibility to manipulate signaling pathways directly in epithelial cells[16-21]. Previous work has established that JAK/STAT signaling is important during tracheal morphogenesis[22-24] and that it regulates epithelial responses to infection and injury in multiple tissues in *Drosophila*[25, 26]. However, the role of this pathway in maintaining airway epithelial homeostasis and coordinating responses to environmental stress has not been systematically investigated.

Here, we use the genetically tractable *Drosophila* airway system to examine how JAK/STAT signaling regulates epithelial homeostasis. We show that basal pathway activity contributes to epithelial integrity, whereas sustained activation is associated with structural remodeling. Furthermore, we demonstrate that environmental stress dynamically modulates pathway activity and that chronic activation induces a coordinated epithelial stress program. Based on these findings, we propose that airway epithelia require a “Goldilocks” level of JAK/STAT signaling, in which balanced pathway activity maintains homeostasis while insufficient or excessive signaling disrupts tissue integrity.

## Results

To investigate how JAK/STAT signaling regulates airway epithelial homeostasis, we combined genetic, environmental, and transcriptomic approaches in the *Drosophila* airway system. We first assessed basal pathway activity and its requirement for epithelial integrity. We then examined how environmental stress modulates pathway activity, followed by analysis of the structural and molecular consequences of sustained pathway activation. Finally, we tested whether pharmacological inhibition can reverse the observed remodeling phenotypes.

### JAK/STAT Signaling Is Present throughout the Larval Tracheal System

To determine whether JAK/STAT signaling is active under physiological conditions, we examined pathway activity using a STAT-responsive transcriptional reporter (Fig. 1A). Under basal conditions, the trachea of third-instar *Drosophila* larvae (L3) displayed near-ubiquitous reporter activity across all tracheal compartments (Fig. 1B-E). However, marked regional differences in signal intensity were evident. Specifically, JAK/STAT signaling was enriched in defined zones of the larval tracheal system, including the dorsal trunks, spiracular branches (SB), and Tr2, regions previously associated with regenerative capacity^26,27^ (Fig. 1D, E). In contrast, tracheal regions fated for apoptosis during metamorphosis (e.g., Tr10) exhibited only minimal reporter activity (Fig. 1C), highlighting a functional correlation between JAK/STAT activity and tissue maintenance.

**Figure 1:**
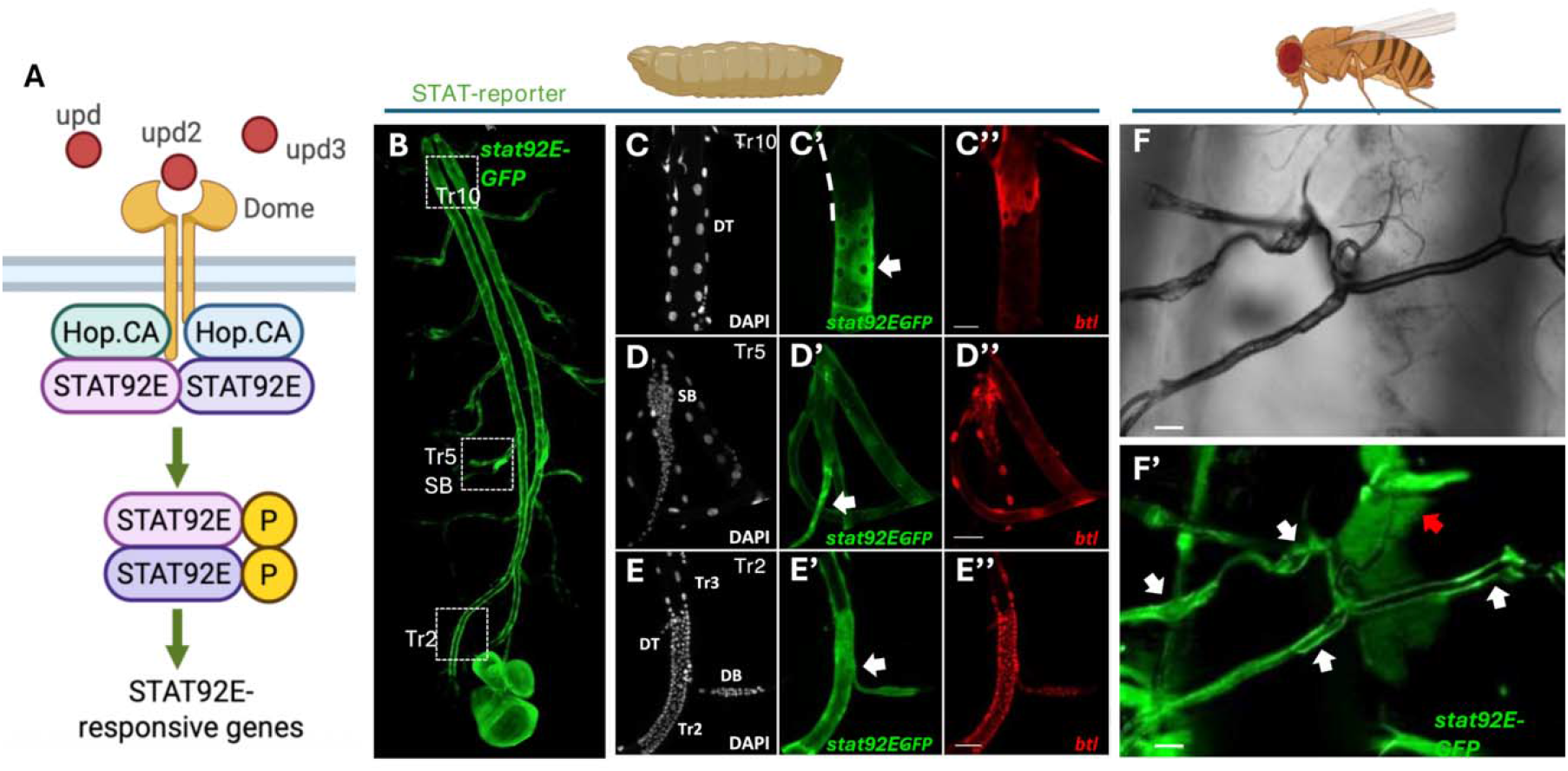
The activation of the JAK/STAT signaling pathway in the respiratory system of *Drosophila*. The activation of the JAK/STAT signaling pathway was detected using a *STAT92E-GFP* reporter (A), which includes *LacZ* expression under control of *btl-Gal4* (*btl-Gal4, STAT92E*-*GFP*; *UAS-LacZ*.*nls)*. (B-E) STAT92E activity pattern (green) in the larval trachea. JAK/STAT activity is present in most tracheal parts (B). Only the posterior part of the trachea (dorsal trunk (DT) of Tr 10, white dashed line in C) showed decreased JAK/STAT signaling activity (and high *btl* expression; red) compared to other tracheal epithelial cells. Conversely, some regions, like the spiracular branch (SB; D, white arrow) and the dorsal branch (DB) in Tr 2 (E, white arrow), showed increased JAK/STAT signaling activity compared to the tracheal epithelial cells with *btl* expression. (F) STAT92E activity pattern in the adult trachea. Scale bar: 50 µm.

In adult flies, reporter analysis revealed sustained pathway activity throughout the airway network (Fig. 1F), confirming a role for JAK/STAT signaling beyond developmental morphogenesis. Transcriptional mapping of *upd2* further localized ligand expression to SB and Tr2 tracheoblasts (Fig. S1), consistent with an epithelial-intrinsic autocrine regulatory circuit.

These observations indicate that JAK/STAT signaling is constitutively active in differentiated airway epithelial cells, suggesting a potential role in maintaining epithelial homeostasis. We next asked whether this basal pathway activity is functionally required for airway epithelial integrity.

### Basal JAK/STAT signaling is required for airway epithelial homeostasis

To assess the functional relevance of basal pathway activity, we reduced pathway activity in airway epithelial cells. We selectively inhibited the pathway in airway epithelial cells using the TARGET system [27] to drive a dominant-negative *Domeless* receptor allele (Dome.DN) (Fig. 2A). Early (embryo) pathway blockade caused larval lethality (Fig. 2B) and led to severe architectural defects, including loss of dorsal trunks and segmental fusion (Fig. 2C). In larval trachea, inhibition of JAK/STAT signaling (here, driven by nach-Gal4) induced epithelial defects characterized by liquid-filled airways, a phenotype that is apparent as a lack of visible trachea in the animals, as liquid filling pushes away the air and thus the differences in refractive indexes between the two media (Fig. 2D, yellow arrows). Moreover, the outcome of the tracheal activation of JAK/STAT signaling critically depended on the time of activation (Fig. S2). At the level of single epithelial cells (here, driven by the vvl-coin system) strong changes in cell structure, reminiscent of apoptosis, occurred (Fig. 2E), which was confirmed by cleaved Dcp-1 staining (Fig. 2F–G).

**Figure 2:**
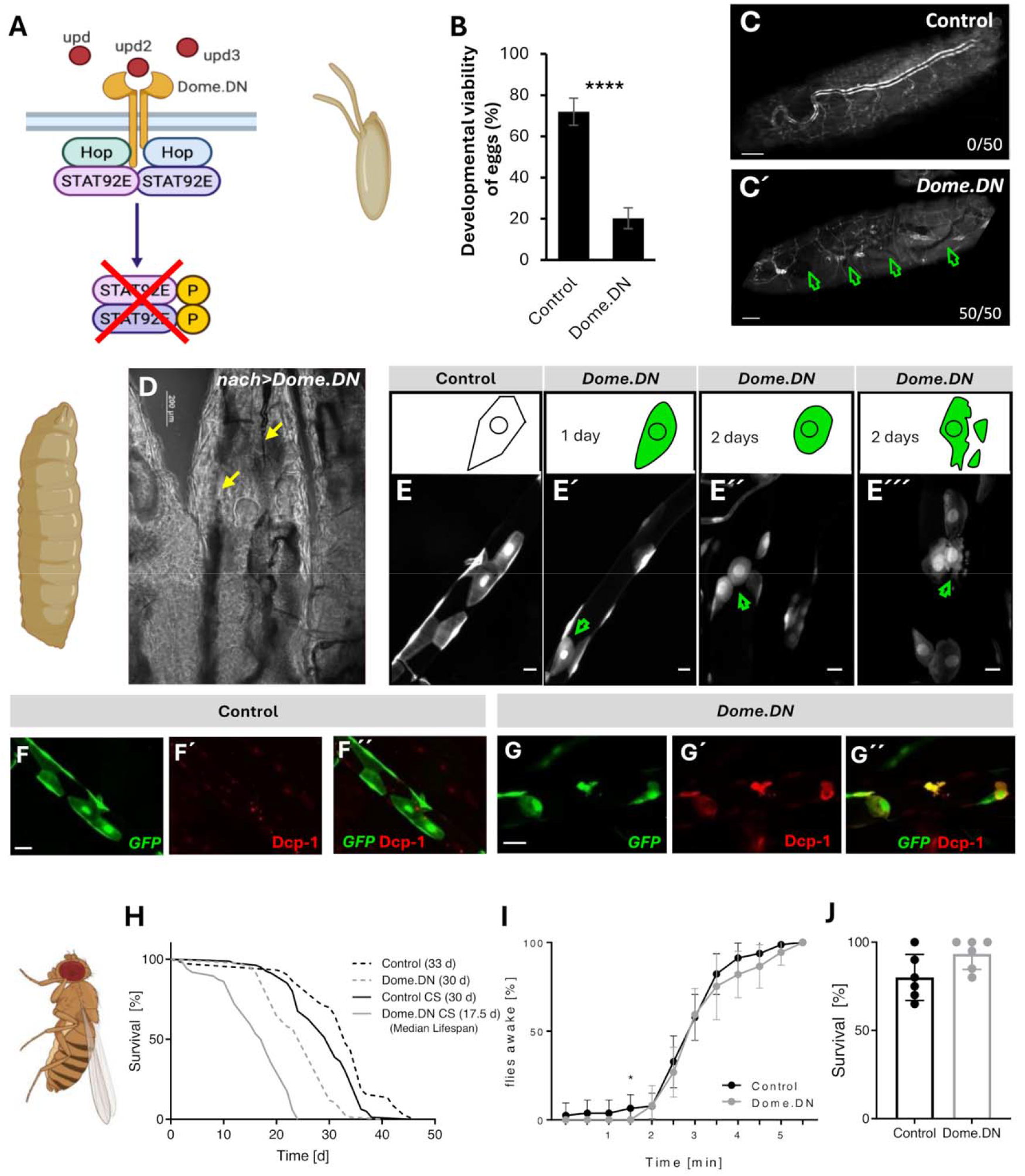
Blocking JAK/STAT signaling induces epithelial cell apoptosis. JAK/STAT signaling was blocked in the trachea by expressing *UAS-Dome*.*DN* under the control of *btl-Gal4;UAS-GFP* (A). (B) The developmental viability in response to JAK/STAT blockade is indicated by the percentage of embryos that hatched. (****p < 0.0001 by Student’s t-test). (C) Blockade of JAK/STAT signaling induced major impairments of the embryonic tracheal structure (arrows). (D) Surviving larvae showed visible defects indicated by liquid-filled regions (arrows). (E) Blocking of the JAK/STAT pathway by expressing *Dome*.*DN* in the trachea led to changes in cell morphology (E’ and E’’, green arrows) and subsequent disintegration (E’’’, green arrow). Scale bar: 20 µm. (F-G) Fluorescence micrographs of the tracheal cells of control *vvl-coin*.*ts* larvae (F), and the tracheal cells of *vvl-coin*.*ts>Dome*.*DN* larvae (G) stained for cleaved Dcp-1 (red, on day 2 after induction). Scale bar: 50 µm. (H) Lifespan analysis under control conditions and cigarette smoke exposure (CS). Median lifespan is indicated in brackets; n=76-103; Log-rank (Mantel-Cox) test. (I, J) Short-term (H; n=6; unpaired t-test) and long-term (I; n=8-9; Mann-Whitney test) hypoxia treatment. * p < 0.05, *** p < 0.001, **** p < 0.0001.

In adults, trachea-specific pathway inhibition reduced organismal lifespan and sensitized animals to chronic CSE exposure, but not to hypoxia (Fig. 2H–J), highlighting stress-specific functional dependencies. Here, CSE and reduced JAK/STAT activity independently reduced lifespan. These findings demonstrate that basal JAK/STAT signaling is required to maintain airway epithelial integrity and organismal stress tolerance. Because airway epithelia constantly encounter environmental challenges, we next investigated whether JAK/STAT signaling responds dynamically to stress.

### Environmental stress activates JAK/STAT signaling in airway epithelial cells

To determine whether environmental stress regulates epithelial JAK/STAT signaling, we exposed animals to several physiologically relevant stress conditions. To evaluate the responsiveness of airway epithelial cells to environmental insults, we employed a GFP-based reporter driven by STAT92E activation in *Drosophila*. Exposure to airborne stressors, including cold air, hypoxia, and cigarette smoke extract (CSE), triggered robust and reproducible pathway activation in larval tracheae (Fig. 3A, B). Similar results were observed in adult flies (Fig. 3C, D). Quantitative imaging confirmed a significant increase in reporter fluorescence under all tested stress conditions. Strikingly, pathway activation co-localized with elevated expression of the cytokine *upd3*, and *upd2* was additionally induced by CSE, whereas other stressors induced much smaller effects (Fig. S3). To test whether a simple intra-organ signaling system is operative, we subjected flies expressing the STAT-reporter, concurrently with RNAi targeting either upd2 or upd3, to hypoxia. Here, we found no rescue of the hypoxia-induced STAT-activation by the RNAi against these most important ligands of the pathway (Fig. 3E, F), which implies that this system is more complex, including other sources of the unpaired ligands or plasticity among different unpaired ligands replacing each other.

**Figure 3:**
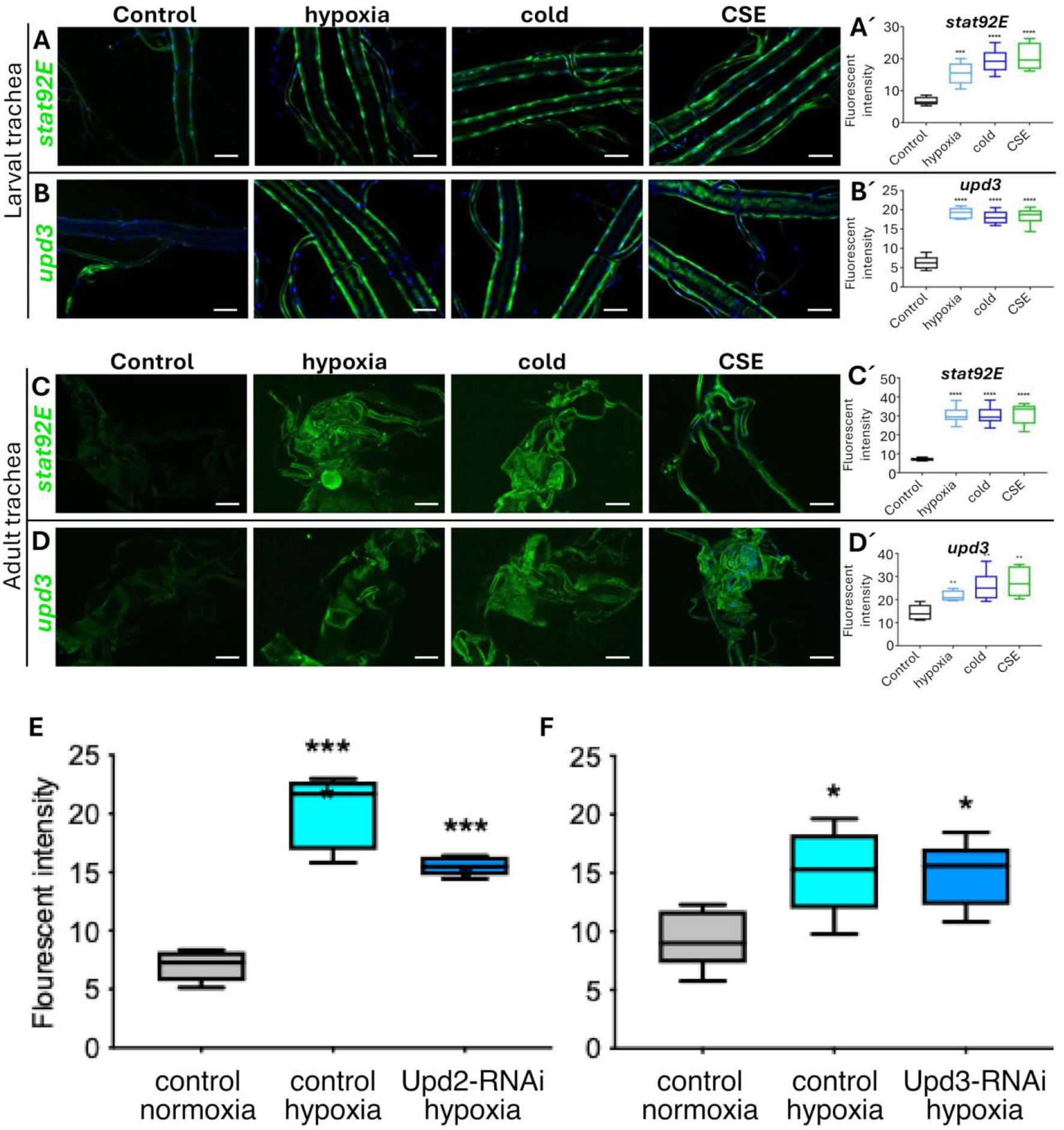
The JAK/STAT signaling pathway in the trachea was activated by external stimuli. (A, C) Fluorescence micrographs of the trachea from *STAT92E*-*GFP* larvae (A) and adults (C) under control conditions and exposed to external stimuli, which included hypoxia, cold air (cold), and cigarette smoke exposure (CSE). The external stimuli induced the activity of STAT92E, which could be observed and quantified through fluorescent intensity (A’ and C’). (B, D) Fluorescence micrographs of the trachea from *upd3*-*Gal4*; *UAS-GFP* larvae (B) and adults (D) under control conditions and exposed to external stimuli. The expression of *upd3* induced by the external stimuli could be observed and quantified through fluorescent intensity (B’ and D’). The fluorescence signal induced by hypoxia in larval trachea was quantified in flies concurrently driving *upd2*-RNAI (E) or *upd3*-RNAi (F). ** *p* < 0.01, *** *p* < 0.001, **** *p* < 0.0001 by Student’s t-test. Scale bar: 100 µm.

These results demonstrate that airway epithelial JAK/STAT signaling functions as a stress-responsive signaling pathway activated by environmental challenges. We next asked how sustained pathway activation affects the structure of the airway epithelium.

### Chronic JAK/STAT activation drives airway epithelial remodeling

To determine the consequences of sustained pathway activation, we chronically activated JAK/STAT signaling in airway epithelial cells. Therefore, we overexpressed either *Hop*.*CA* (constitutively active JAK) or the cytokine *upd3* (Fig. 4A). Both perturbations induced at the embryonic state led to embryonic or early larval lethality (Fig. 4B, C). The structural analysis of these perturbations revealed severe epithelial disruption. Here, *upd3* overexpression yielded more severe epithelial disruption than *Hop*.*CA* overexpression (Fig. 4D-E). Conditional activation of the pathway in larvae via the TARGET system resulted in structural changes in the tracheal system (Fig. 4F, G). Here, structural defects became apparent, including airway stenoses. A closer analysis of the structural changes of the dorsal trunks of L3 larvae using a mosaic system based vvl-coin revealed a narrowing of the airway lumen only in the affected epithelial cells (Fig. 4F, G). A detailed analysis of the chitinous intima of the larval trachea showed that the usually highly regular taenidial structure was strongly impaired and characterized by irregularities (Fig. S4).

**Figure 4:**
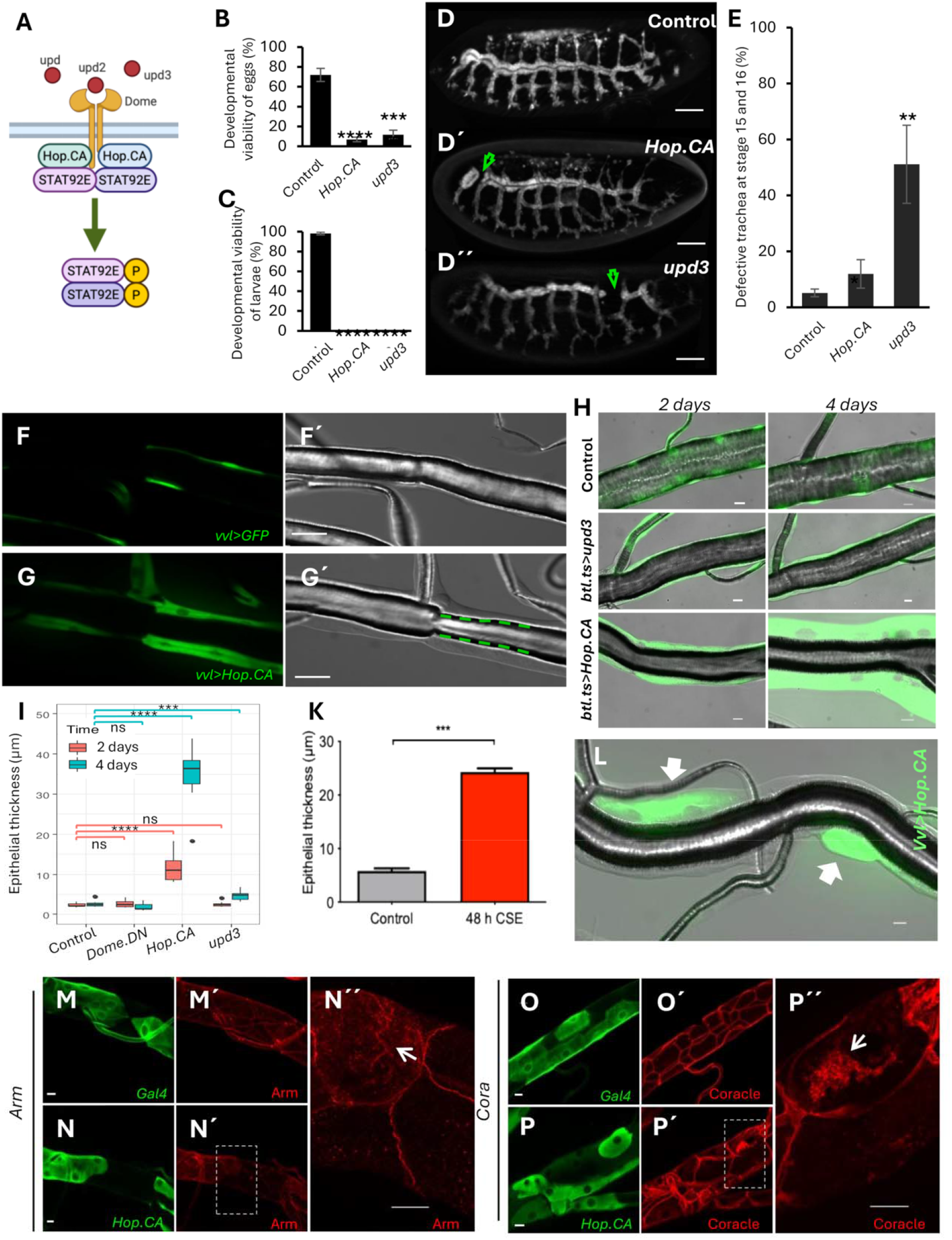
Ectopic JAK/STAT signaling activation in the trachea induced structural changes. Ectopic activation (*Hop*.*CA* or *upd3*) of epithelial JAK/STAT signaling was achieved by ectopic activation of the pathway (A). (B, C) Signaling pathway activation during development (*btl>Hop*.*CA* or *btl*>*upd3*) induced lethality. The percentage of embryos that hatched (B) and the percentage of larvae that developed into pupae (C) are shown. (D-E) Incomplete tracheal development during tracheal morphogenesis. (F, G) Micrographs of DT8 of *vvl-FLP, CoinFLP-Gal4, UAS-EGFP* (*vvl-coin*)*>Hop*.*CA* larvae. Scale bar: 20 µm. The thickening of the tracheal epithelium is observed in those cells that express *Hop*.*CA* (arrows). A narrowing of the airspace was observed in the affected epithelial cells (G; dotted lines indicate the airway boundaries), whereas unaffected cells remained inconspicuous (F). (H) Ectopic activation of the pathway either by upd3 or Hop.CA overexpression for 2 days or 4 days using the Target system by switching the temperature in the larval stage from 19°C to 30°C. Epithelial cells express GFP. Scale bar: 20 µm. (I) Quantification of the epithelial thickness (DT8) in response to upd3-, Hop.CA-, or Dome.DN overexpression for different times. (K) Quantitative evaluation of epithelial thicknesses of affected regions of the trachea following repeated chronic smoke exposure. (L) Evaluation of the thickening phenotype using the mosaic-inducing vvl system (green are affected cells), unaffected cells remained normal. (M-P) Localization of the membrane-associated proteins armadillo (arm) and Coracle (cora) in control epithelial cells and those showing strong JAK/STAT activation. (M, N) Armadillo staining showed accumulation in the cell of the immunoreactivity (arrow). (O, P) Similar localization errors are shown for Coracle staining (arrow). ns = not significant, * *p* < 0.05, ** *p* < 0.01, *** *p* < 0.001, **** *p* < 0.0001; Student’s t-test.

These findings indicate that excessive JAK/STAT signaling induces pathological airway remodeling, suggesting that epithelial homeostasis requires tightly controlled pathway activity. We next sought to determine the cellular mechanisms underlying this remodeling phenotype.

### JAK/STAT hyperactivation alters cell structure and junctional organization

In response to JAK/STAT activation, the tracheal epithelial cells exhibited pronounced epithelial thickening. This thickening response was observed in response to *Hop*.*CA* and *upd3* overexpression, with much higher thickening in response to *Hop*.*CA* (Fig. 4H, I). In addition to the other structural defects observed, the localization of junctional proteins and epithelial polarity markers was analyzed in control and *Hop*.*CA*-expressing airway epithelial cells. This phenotype was fully cell-autonomous, as demonstrated by mosaic analysis (Fig. 4L), and driven by increased cell volume rather than hyperplasia (Fig. S5). We observed locally occurring thickening of the epithelium to strong CSE application (Fig. 4K), resembling reactions to infection in the larval airway epithelium [28]. Furthermore, weaker expression drivers (*nach*-Gal4) produced the same phenotype with reduced severity (Fig. S6). Epithelial thickening in the trachea also occurs in response to activation of the IMD pathway (TNF-α receptor equivalent, induced by *btl-Gal4, UAS-PGRP-LC*) [29]. To test, if JAK/STAT pathway activation uses the same mechanisms, we blocked STAT expression while activating the IMD pathway and found no phenotype rescue (Fig. S6E).

To investigate cellular mechanisms underlying epithelial remodeling, we examined the localization of junctional proteins (Coracle (Cora) and Armadillo (Arm)). In control tissues, these proteins were restricted to cell–cell boundaries. In contrast, sustained JAK/STAT activation was associated with intracellular accumulation of junctional components, suggesting altered protein trafficking or localization. Using mosaic analysis, we observed intracellular accumulation of Arm and Cora, respectively, in affected cells (Fig. 4M-P), which is not seen in control cells. To investigate the molecular programs underlying these structural changes, we next performed transcriptomic analysis of airway epithelial cells with activated JAK/STAT signaling.

### Sustained JAK/STAT Activation induces a Coordinated Transcriptional Response

Transcriptomic analysis of airway epithelial cells with sustained JAK/STAT activation identified widespread changes in gene expression. RNAseq data are deposited in the GEO database (GSE212132). Upregulated genes included canonical pathway targets as well as genes associated with immune responses, stress pathways, and metabolism. Downregulated genes were enriched for functions related to tissue structure and differentiation (Fig. 5A–B). A closer analysis revealed major upregulated genes (Table 1) and the KEGG pathways involved (Table S1). Transcription factor binding site analyses revealed STAT92e as one of the hits (Table 2).

**Table 1:** Genes that were highly upregulated in response to *Hop*.*CA* overexpression.

| Name | Maximum group mean | Fold change | FDR p-value |
| --- | --- | --- | --- |
| CG16772 | 11.85 | 135.44 | 0 |
| hop | 286.06 | 35.43 | 0 |
| CG14570 | 42.21 | 35.37 | 0 |
| CG14147 | 55.31 | 32.88 | 0 |
| Hsp70Bb | 161.51 | 29.99 | 0 |
| Ubx | 200.46 | 23.51 | 0 |
| pre-mod(mdg4)-I | 1.12 | 18.67 | 2.22648E-06 |
| IM14 | 11.46 | 15.14 | 3.80132E-07 |
| IM4 | 26.43 | 14.97 | 1.74645E-12 |
| CG14569 | 1576.88 | 14.73 | 0 |
| IM2 | 7.26 | 14.47 | 2.01792E-06 |
| CG5791 | 1.79 | 13.80 | 2.90502E-05 |
| CG9121 | 3.41 | 11.54 | 0 |
| IM1 | 7.16 | 10.97 | 7.52122E-05 |
| CG11413 | 35.70 | 10.64 | 3.44242E-07 |
| CG16710 | 2.13 | 10.32 | 1.72352E-07 |
| CG31960 | 267.26 | 10.05 | 0 |
| CG16789 | 1.45 | 9.46 | 5.58576E-09 |
| CG6034 | 6.92 | 8.47 | 4.7943E-11 |
| Ace | 2.06 | 8.19 | 0 |
| CG13059 | 9.90 | 7.58 | 1.33893E-06 |
| CG31809 | 8.64 | 7.39 | 0 |
| CG7203 | 9.41 | 7.21 | 8.66989E-09 |
| Vha14-2 | 2.51 | 7.05 | 2.31024E-07 |
| wb | 7.35 | 6.69 | 0 |
| Drs | 8166.47 | 6.64 | 0 |
| CG5778 | 3.20 | 6.61 | 0.000294835 |
| CG3457 | 2.69 | 6.34 | 2.38755E-06 |
| Lcp65Ad | 9.93 | 5.79 | 2.58249E-08 |
| lectin-22C | 29.06 | 5.67 | 6.88538E-06 |
| fuss | 3.97 | 5.67 | 0 |
| CG43333 | 2.39 | 5.63 | 1.48392E-09 |
| CG18336 | 2.96 | 5.49 | 5.52167E-05 |
| CG32563 | 3.06 | 5.35 | 0.000696584 |
| FASN2 | 1.65 | 5.34 | 2.95545E-10 |
| Cyp4d21 | 45.70 | 5.31 | 0 |
| Obp57d | 1.80 | 5.28 | 0.000102926 |
| CG17560 | 2.31 | 5.22 | 0.000766701 |
| CG5888 | 68.16 | 5.20 | 0 |

$p < 1 \times 10^{-4}$ , fold change > 5, maximum group mean > 1. The list is restricted to 39 genes.

**Table 2:** Transcription factor-binding site motifs enriched in 41 highly upregulated genes.

| Matrix ID | Matrix name | P-value |
| --- | --- | --- |
| MA0023.1 | dl (var.2) | 0.000443261 |
| MA0022.1 | dl | 0.00407113 |
| MA0532.1 | STAT92E | 0.00794732 |
| MA0450.1 | hkb | 0.00811969 |
| MA0197.2 | nub | 0.0154201 |
| MA0242.1 | Bgb::run | 0.0371696 |
| MA0204.1 | Six4 | 0.0443521 |
| MA0444.1 | CG34031 | 0.0488169 |

A total of 133 transcription factor profiles were used.

**Figure 5:**
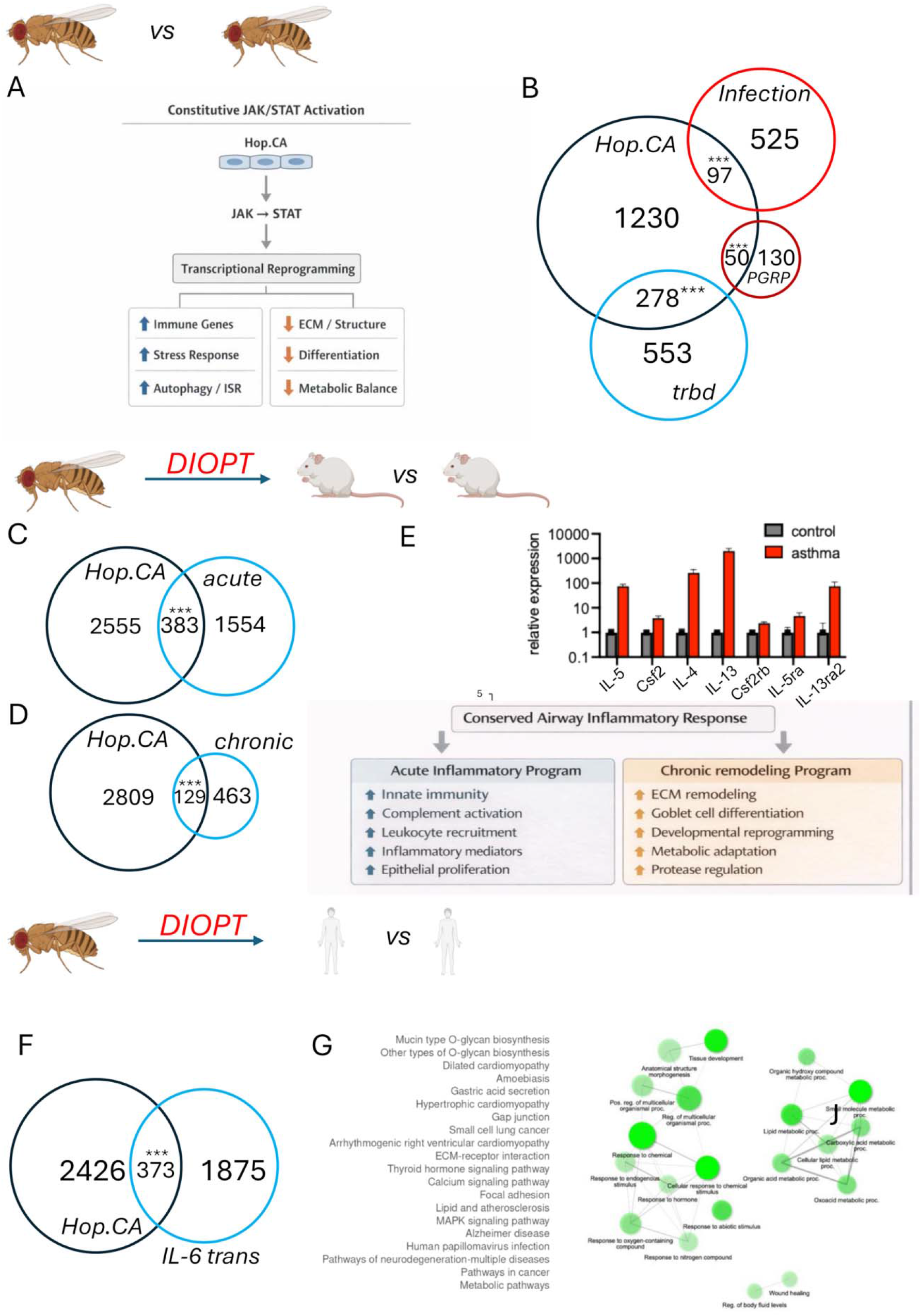
Analysis of genes differentially expressed between control trachea and those ectopically activated by Hop.CA overexpression. (A) Schematic overview of the DEGs that are up- or downregulated in response to Hop.CA expression. (B) Venn diagram showing DEGs in response to Hop.CA overexpression compared with those following Trabid-KO, tracheal infection, or ectopic activation of the IMD pathway (PGRP-LC overexpression). (C) Using DIOPT to identify mouse orthologs of the Hop.CA DEGs from (A), these orthologs were compared with DEGs from lungs subjected to acute and chronic asthma models. (D) Key chemokines and chemokine receptors induced by OVA aerosol are involved in the JAK/STAT signaling pathway. (E) Analysis of the genes commonly regulated between Hop. CA-induced genes and those from the acute and chronic asthma models. (F) DIOPT-assisted identification of transformation of Hop.CA DEGs into human orthologs, which were compared with genes differentially expressed in human airway epithelial cells exposed to IL-6 trans-signaling. The common genes were analyzed regarding GO categories in the biological processes group and through a string analysis of these data (G). *** *p* < 001; Fisher’s exact test.

Comparison with previously described gene sets revealed partial overlap with immune and stress-related transcriptional programs (Trabid-deficiency, tracheal infection, and PGRP-LE overexpression signatures [29-31]). These findings suggest that JAK/STAT activation induces a composite transcriptional state integrating multiple regulatory pathways. The corrected Trabid gene set (n = 890) showed a substantial overlap of 291 genes with the Hop dataset (32.7% of the Trabid set; Jaccard index = 0.113). Hypergeometric testing revealed a >2-fold enrichment over random expectation (expected ≈127 genes; observed = 291; p < 10^−100^), indicating strong biological convergence. The overlapping genes included canonical antimicrobial peptides (e.g., *Drs, IM4*), heat shock proteins (*Hsp22, Hsp70Bb*), cytochrome P450 enzymes, and lipid metabolic regulators, consistent with activation of immune, stress, and metabolic remodeling programs. Comparisons with Infection and PGRP-LE signatures revealed partial but not complete overlap, supporting the interpretation that Hop activation induces an infection-like state that extends beyond classical Imd pathway activation.

Together, these findings demonstrate that Hop signaling engages a composite transcriptional program integrating immune activation, proteotoxic stress responses, and metabolic reprogramming.

### Cross-Species Comparison Reveals Conserved Features of Epithelial Stress Responses

To assess the broader relevance of these transcriptional changes, we compared the *Drosophila* dataset with gene expression profiles from murine asthma models and cytokine-stimulated human airway epithelia. These analyses revealed shared features, including activation of immune signaling, oxidative stress responses, and alterations in epithelial transport and metabolism. To enable the direct comparison, we generated lists of mouse homologs of the regulated *Drosophila* genes using the DIOPT online tool [32]. The overlaps are highly significant (Fig. 5C, D), with more than 15-20% overlap with the mouse data. Constitutive Hop/JAK signaling induces a transcriptional program characterized by activation of innate immune effectors, antimicrobial peptides, stress responses, and secretory pathway components, accompanied by extensive remodeling of metabolism and trafficking. We evaluated this murine dataset by testing cytokine genes and selected genes involved in vesicular transport by qRT-PCR (Fig. 5E). The direct comparison with the acute mouse asthma model revealed a common signature for the upregulated genes, including 1) antimicrobial/innate immune activation, 2) extensive cytokine signaling, 3) oxidative stress response, and 4) increased secretory activity. Downregulated are especially programs associated with tissue differentiation and metabolic homeostasis (Fig. 5F).

Furthermore, we compared the RNAseq data with those derived from human airways subjected to a strong and specific JAK/STAT stimulus, namely IL-6 transsignaling [33]. Again, we observed an overlap in the DEGs with the *Drosophila* data transformed using the DIOPT pipeline (Fig. 5G). Commonalities of the differentially expressed genes comprise a coordinated transcriptional program characterized by activation of ER stress, inflammatory signaling, epithelial transport, and metabolic pathways. Genes involved in the unfolded protein response and ER proteostasis (*HSPA5, HSP90B1, HYOU1, ATF4, ERP44*) and ER–Golgi trafficking (*SEC23B, SEC31B, COPB1*) were enriched. Concurrently, key components of innate immune signaling and NF-κB activation (*TLR1, IRAK1, IRAK4, RELA, RELB, BIRC3*) and oxidative stress regulators (*NOX1, DUOX1, DUOX2, PRDX4, CAT*) were detected. The gene set also included epithelial transport and barrier-associated genes such as *CFTR, SCNN1A/B/G, SLC26A9*, and *AQP3/AQP4*, together with extracellular matrix and adhesion genes (*FBLN1, EFEMP1, LAMC2, ITGA family*). Additionally, enzymes of the mevalonate and lipid metabolism pathways (*HMGCR, HMGCS1, MVD, PLIN2/3/4/5*) were enriched, indicating coordinated metabolic remodeling (Fig. 5H, I).

While these comparisons are based on orthology mapping and do not imply direct equivalence, they suggest that key aspects of epithelial stress responses may be conserved across species. Finally, we asked whether the airway remodeling phenotype can be rescued by pharmacological intervention.

### Pharmacological Inhibition of JAK/STAT Reverses Epithelial Remodeling

To determine whether airway remodeling depends on sustained pathway activation, we tested whether pharmacological inhibition of JAK signaling could reverse epithelial defects. Treatment of *Hop*.*CA*-expressing flies with a JAK inhibitors with a JAK1-like profile (Oclacitinib, Baricitinib, and Filgotinib) reduced epithelial thickening. Other JAK inhibitors with a different pharmacological profile (Tofacitinib, CYT387, Ruxolitinib) failed to rescue the phenotype (Fig. 6A).

**Figure 6:**
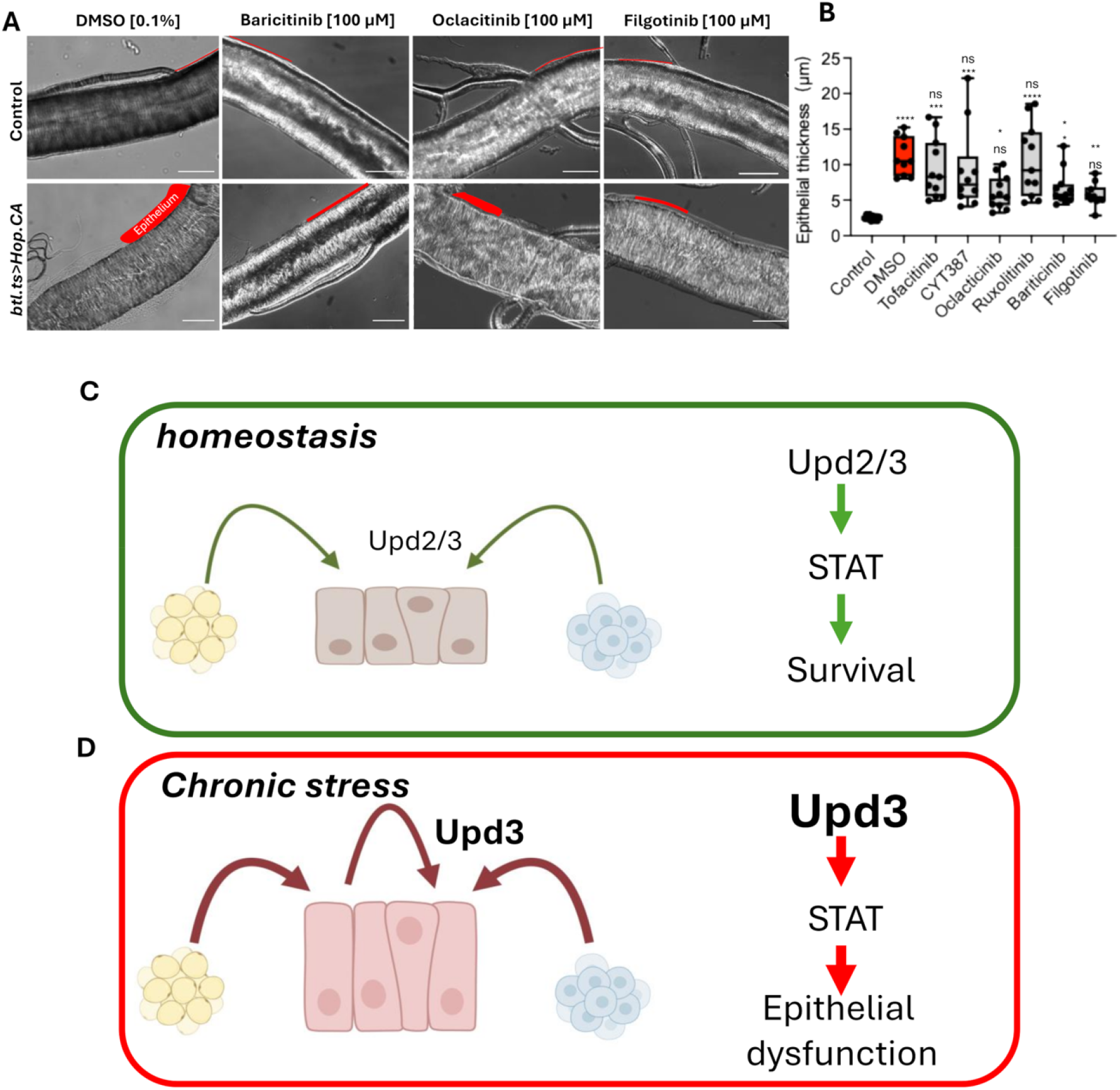
Pharmacological rescue of the epithelial thickening phenotype and concluding model of the JAK/STAT action in airway epithelial cells. (A, B) Changes in tracheal epithelial thickness could be rescued by applying specific JAK inhibitors. (A) Microscopy images of the tracheal epithelium (3^rd^ instar larvae) of Control (*btl*.*ts>w*^1118^) compared to activated JAK/STAT signaling pathway (*btl*.*ts>Hop*.*CA*). Scale bar: 50 µm. (B) Quantification of epithelial thickness (highlighted in red) in JAK inhibitor treatment groups compared to the DMSO control. ** *p* < 0. 01; Student’ s t-test. (C) Schematic characterization of JAK/STAT signaling under homeostatic conditions and in response to chronic stress leading to or prolonged overactivation (D).

### Summary

Together, these results demonstrate that airway epithelial homeostasis requires JAK/STAT signaling to operate within a narrow functional range. Basal signaling maintains epithelial integrity and stress resistance, whereas chronic hyperactivation induces pathological airway remodeling through conserved epithelial stress programs (Fig. 6C, D). Together, these findings demonstrate that airway epithelial homeostasis requires JAK/STAT signaling to operate within a narrow functional range. Basal pathway activity supports epithelial integrity and stress resistance, whereas sustained hyperactivation disrupts epithelial architecture and induces conserved airway remodeling programs.

## Discussion

### Basal Homeostasis and Epithelial Viability

Leveraging the genetically streamlined architecture of *Drosophila melanogaster*, this study uncovers the bifunctional role of the JAK/STAT cascade in airway epithelia. Because the system relies on a single receptor (Domeless), kinase (Hopscotch), and transcription factor (STAT92E) [34], we circumvented vertebrate redundancy [35] to dissect pathway dynamics at high resolution. Our data indicate that JAK/STAT signaling operates within a narrow functional range in airway epithelial cells. Basal pathway activity is required to maintain epithelial integrity and organismal stress resistance, whereas sustained hyperactivation induces structural remodeling and disrupts epithelial organization. Together, these findings support a “Goldilocks” model in which balanced JAK/STAT signaling preserves airway homeostasis, while both insufficient and excessive activity impairs epithelial function.

This requirement for basal activity perfectly aligns with recent mammalian findings demonstrating that tonic JAK/STAT signaling is critical for maintaining cellular homeostasis and identity across immune and epithelial compartments [36, 37]. Furthermore, our observation that early pathway disruption causes catastrophic architectural defects during tracheal tube formation echoes vertebrate studies establishing STAT3 as a fundamental requirement for lung organogenesis and epithelial lineage maintenance [12, 38].

### Stress Responsiveness and Dynamic Signaling

The airway epithelium represents the primary interface between the organism and environmental stressors [39]. In this study, exposure to hypoxia, cold air, and cigarette smoke extract (CSE) induced robust activation of JAK/STAT signaling in airway epithelial cells. This response was associated with increased expression of Upd ligands, suggesting that epithelial cells actively participate in amplifying stress signals, but that they are not the only source of upd expression in response to stressors.

These findings indicate that JAK/STAT signaling functions as a stress-responsive pathway in airway epithelia. Importantly, our data suggest that both insufficient and excessive pathway activity are detrimental: reduced signaling sensitizes the organism to stress, whereas sustained activation is associated with structural remodeling. Together, this supports a model in which airway epithelial homeostasis depends on maintaining JAK/STAT signaling within a defined functional range.

### A Goldilocks Principle of JAK/STAT Signaling in Airway Epithelia

Our findings support a “Goldilocks” principle of JAK/STAT signaling in airway epithelia, in which both insufficient and excessive pathway activity compromise tissue integrity. Basal signaling appears to support epithelial survival and stress resistance, whereas chronic hyperactivation is associated with epithelial remodeling and functional disruption.

This conceptual framework provides a unifying explanation for the dual roles of JAK/STAT signaling observed in both physiological and pathological contexts. While acute activation likely supports adaptive responses, sustained signaling may shift the system toward a maladaptive state. This echoes the delicate balance of the JAK/STAT pathway in airway epithelia [12]. If the pathway fails to activate, the organism is heavily sensitized to chronic stress. Conversely, as seen in chronic lung conditions like asthma and COPD where the JAK/STAT pathway remains persistently hyperactive [10, 40], the protective stress response transforms into pathology.

### Stress-Induced Transcriptional Reprogramming

Transcriptomic profiling of airway epithelial cells with sustained JAK/STAT activation revealed a coordinated transcriptional response characterized by upregulation of innate immune effectors, stress response pathways, and metabolic regulators. These findings suggest that chronic pathway activation induces a broad epithelial stress program rather than a purely immune-specific response.

Comparison with previously described gene sets, including immune and stress-related signatures, revealed significant overlap, indicating that JAK/STAT activation engages conserved regulatory modules [30, 31]. The transcriptional reprogramming induced by sustained JAK/STAT activation seems to fundamentally alter the cellular identity, shifting it into a systemic, stress-adapted state that closely mimics the inflammatory landscapes seen in idiopathic pulmonary fibrosis and acute lung injury [41, 42]. However, these overlaps are based on orthology mapping and should be interpreted cautiously, as species-specific differences in gene regulation are not fully captured.

### Molecular Mechanisms Underlying Airway Remodeling

Our data support a model in which sustained JAK/STAT activation induces a coordinated stress response that increases secretory demand and proteostatic load within epithelial cells. This imbalance is likely to affect vesicular trafficking and protein localization, as suggested by the intracellular accumulation of junctional proteins such as Coracle and Armadillo.

These observations are consistent with impaired trafficking and junctional organization contributing to epithelial thickening and lumen narrowing. This illustrates how the transcriptomic shift toward increased, yet defective, secretory activity compromises the delivery of critical structural components, ultimately leading to barrier failure and organismal lethality, a structural decline reminiscent to the epithelial-to-mesenchymal transition (EMT) seen in progressive lung disease [43]. While the present study does not establish a direct causal sequence between these processes, the data support a mechanistic link between sustained pathway activation, cellular stress responses, and structural remodeling.

### Conservation Across Species and Disease Contexts

Cross-species comparisons revealed overlap between the transcriptional signatures induced by JAK/STAT activation in *Drosophila* and those observed in murine asthma models and cytokine-stimulated human airway epithelia. These shared features include activation of immune signaling pathways, stress responses, and alterations in epithelial transport and metabolism.

While these comparisons rely on computational mapping and do not imply direct equivalence, they suggest that core aspects of epithelial stress responses may be conserved across species. This supports the use of the *Drosophila* airway system as a model to investigate general principles of epithelial remodeling.

### Pharmacological Modulation and Translational Implications

Pharmacological inhibition experiments demonstrated that airway remodeling induced by sustained JAK/STAT activation can be partially reversed in vivo. Notably, inhibitors with higher specificity toward JAK1-like activity showed greater efficacy in reducing epithelial thickening. These results precisely mirror the specific kinase binding profiles established in vertebrate models [44-46]. This functional divergence suggests that the hyperactive *Drosophila* Hop.CA kinase shares a profound structural or functional homology with vertebrate JAK1, establishing this in vivo model as a highly specific screening platform for next-generation, JAK1-targeted respiratory therapeutics [16, 17, 47]. Although inhibition of the JAK/STAT signaling pathway is currently being actively explored as a therapeutic strategy, particularly for chronic forms of asthma that are refractory to conventional treatments [48], it may interfere with the regenerative capacity of the airway epithelium [13]. Thus, therapeutic strategies will likely require careful modulation rather than complete inhibition.

## Conclusion

In summary, our findings define a narrow functional window for JAK/STAT signaling in airway epithelia, in which balanced pathway activity is required to maintain epithelial integrity and stress resilience. Deviations from this balance—either through insufficient or excessive signaling—lead to epithelial dysfunction and structural remodeling. This “Goldilocks” principle provides a conceptual framework for understanding how epithelial tissues integrate environmental signals while preserving homeostasis and offers insights into conserved mechanisms underlying airway disease.

## Materials and Methods

### Drosophila strains and husbandry

*STAT92E-GFP* was used to monitor the pathway’s activation [49]; the Gal4-UAS system [50] targeted ectopic expression to the tracheal system. The Gal4 drivers included: *btl-Gal4, UAS-GFP* on the 3rd chromosome, and *btl-Gal4, UAS-GFP* on the 2nd chromosome (obtained from the Leptin group, Heidelberg, Germany); *upd2-Gal4*; *upd3-Gal4 [19]*; *nach-Gal4 [51]*. The UAS responders consisted of: *UAS-LacZ*.*nls* (BDSC 3956), *UAS-domeΔcyt2*.*1* (*Dome*.*DN*), and *UAS-Hop*.*CA* (*UAS-hopTumL*), which were obtained from N. Perrimon [22, 52]. The *UAS-upd3* was constructed in our lab. *TubP-Gal80[ts]* (BDSC 7018) was obtained from the Bloomington stock center. Unless otherwise stated, the flies were raised on standard medium at 25 °C with 50–60% relative humidity under a 12:12□h light/dark cycle as described earlier [53].

### Coin-FLP expression system

*Vvl-FLP/CyO; btl-moe*.*mRFP* (BDSC 64233), *tubP-Gal80[ts]* and *CoinFLP-Gal4, UAS-2xEGFP* (BDSC 58751) were used to construct animals for the tracheal mosaic analysis. Ventral veins lacking (*vvl*) was expressed in larval tracheal clones that covered approximately 30 to 80% of the trachea [54, 55]. The genotype of the flies was *vvl-FLP, CoinFLP-Gal4, UAS-2xEGFP*/*CyO* (*vvl-coin*) and *vvl-FLP, CoinFLP-Gal4, UAS-2xEGFP*/*CyO*; *tub-Gal80[ts]* (*vvl-coin*.*ts*).

### Developmental viability

For the developmental viability of eggs, they were collected overnight and not physically handled in any way. The number of hatched eggs and pupae were counted starting two days after collection. Each group had four replicates that included more than 20 eggs. Tub-Gal80 [ts] was used to limit UAS responder expression during the larval stage to study the developmental viability of larvae. Animals were raised at 18 °C to keep the UAS responder gene silent. Larvae at different instar stages were transferred to a new medium at 29 °C. In this study, four replicates were typically performed with 30 larvae each. The stage of the larvae was determined by the appearance of the anterior spiracles.

### Determination of epithelial thickness

The L2 and L3 larvae trachea were carefully dissected from the posterior side of the body in PBS. The isolated trachea were immersed in 50% glycerol, and digital images were captured within 15 min. L2 larvae were distinguished from L3 larvae by the appearance of anterior spiracles. The relative ages of L3 larvae were inferred from the animal’s size. Thirty larvae were used in each group, and specimens were analyzed by Image J.

### Drug application in Drosophila

JAK inhibitors (Baricitinib #16707, Oclacitinib #18722, Filgotinib #17669 - Cayman Chemicals, Michigan, USA) were diluted in DMSO [100 mM]. For later application, the inhibitors were diluted 1:10 in 100% EtOH. We used 20 µl of each diluted inhibitor for 2 ml of concentrated medium (5% yeast extract, 5% corn flour, 5% sucrose, 1% low-melt agarose, 1 ml of 10% propionic acid, and 3 ml of 10% Nipagin). The eggs of each crossing have been applied to the modified medium and kept at 20 °C until the larvae reach the L2 stage. Afterwards, they were incubated at 30 °C for 2 days to induce the *btl-Gal4, UAS-GFP; tub-Gal80[ts]* (*btl*.*ts*)–driver. The trachea of L3 larvae were carefully dissected from the posterior side of the body in PBS. The isolated trachea was immersed in 50% glycerol, and digital images were captured every 15 minutes. 10 larvae were used for each group.

### Experimental design for hypersensitivity pneumonitis in mice

Experimental design for hypersensitivity pneumonitis contains two processes, sensitization and challenge of mice with ovalbumin (Sigma Aldrich, A5503). The protocol refers to the protocol that [56] is described with a few modifications. 8-week-old Balb/C mice were sensitized with intraperitoneal injections of 20 µg of OVA emulsified in aluminum hydroxide in a total volume of 1 ml on days 7 and 14, followed by three consecutive challenges each day by exposure to OVA or PBS aerosol for 30 min. Mice were sacrificed 24 h following the final challenge. The left lungs were collected for histological analysis, and the superior lobes were dissected for RNA analysis. Six mice per group were used in this experiment and four mice were randomly selected in each group for analyses. Genesky performed total mRNA sequencing, data processing, and statistical analysis (Shanghai, China). The experimental protocols were approved by the Animal Care and Protection Committee of Weifang Medical University (2021SDL418).

### AB-PAS staining and immunofluorescence analysis of mouse lung

Mice were sacrificed in an excess of CO_2_ gas. The lungs of euthanized mice were inflated by intratracheal injection of cold 4% paraformaldehyde and then fixed and embedded in paraffin as described in [57]. AB-PAS staining and Immunofluorescence analysis were supported by Servicebio (Wuhan, China). All sections were photographed using a microscope slide scanner (Pannoramic MIDI: 3Dhistech). The materials and methods could be found at the following link https://www.servicebio.cn/data-detail?id=3040&code=MYYGSYBG and https://www.servicebio.cn/data-detail?id=3595&code=RSSYBG. We list the primary reagents here: AB-PAS solution set (Servicebio, G1049), Anti-CD11b (Servicebio, GB11058), and DAPI (Servicebio, G1012-10ML).

### Time-lapse microscopy

All images were acquired using a ZEISS Axio Image Z1 and a ZEISS LSM 880 fluorescent microscope (INST 257/591-1 FUGG). Embryos were dechorionated in 3% sodium hypochlorite and immersed in Halocarbon oil 700 (Sigma Aldrich, 9002-83-9, Deisenhofen, Germany). Then the embryos were imaged after stage 15 when the tracheal tree formed at 3-hour intervals. 40 embryos were investigated in each group.

### Cigarette smoke and hypoxic exposure

All cigarette smoke exposure experiments were carried out in a smoking chamber, attached to a diaphragm pump. Common research 3R4F cigarettes (CTRP, Kentucky University, Lexington, USA) were used for all experiments. The vials containing animals were capped with a monitoring grid to allow the cigarette smoke to diffuse into the vial. For long-term smoke experiments, L2 larvae were exposed to smoke thrice a day for 30 minutes each, on two consecutive days. For heavy smoke experiments, L3 larvae were exposed to the smoke of 2 cigarettes for 45 min, which led to about 35-50% mortality. To study the effects of hypoxia on the activity of JAK/STAT signaling pathway, larvae were exposed to long-term and short-term hypoxia separately. For long-term hypoxia experiments, L2 larvae were exposed to 5% oxygen three times a day for 30 min each, on two consecutive days. L3 larvae were exposed to 1% oxygen once for 5 hours for short-term hypoxia experiments. At least 20 animals in 4 vials were investigated in each group. Cigarette smoke and hypoxia exposure of adults was performed as described earlier [19] .

### Immunohistochemistry of Drosophila trachea

Larvae were dissected by ventral filleting and fixed in 4% paraformaldehyde for 30 min. Embryos were staged according to Campos-Ortega and Hartenstein [58] and fixed in 4% formaldehyde for 30 min. Immunostaining was performed following standard protocols as described earlier[59, 60]. GFP signals were amplified by immunostaining with polyclonal rabbit anti-GFP (used at 1:500, Sigma-Aldrich, Merck KGaA, Darmstadt, Germany, SAB4301138). 40-1a (used at 1:50, DSHB, Iowa City, USA) was used to detect Beta-galactosidase. Coracle protein was detected with a monoclonal mouse anti-coracle antibody (used at 1:200, DSHB, Iowa City, USA, C566.9). Armadillo protein was detected with a monoclonal mouse anti-armadillo antibody (DSHB, US, N2 7A1, used at 1:500). A monoclonal rabbit Cleaved Drosophila Dcp1 (used at 1:200, Cell Signaling, Frankfurt/M, Germany, #9578) was used to detect apoptotic cells. Secondary antibodies used were: Cy3-conjugated goat-anti-mouse, Cy3-conjugated goat-anti-rabbit, Alexa488-conjugated goat-anti-mouse (used at 1:500, Jackson Immunoresearch, Dianova, Hamburg, Germany), Alexa488-conjugated goat-anti-rabbit (used at 1:500, Cell signaling, Frankfurt/M, Germany, #9578). Tracheal chitin was stained with 505 star conjugated chitin-binding probe (NEB, Frankfurt/M, Germany, used at 1:300). Nuclei were stained with 4’,6-Diamidino-2-Phenylindole, Dihydrochloride (DAPI) (Roth, Karlsruhe, Germany, 6843). 30 specimens were investigated in each group. Specimens were analyzed and digital images were captured using either a confocal microscope (Zeiss LSM 880, Oberkochen, Germany) or a conventional fluorescence microscope (ZEISS Axio Imager Z1, Zeiss, Oberkochen, Germany).

### RNA isolation and RNA sequencing

For the gene expression analysis of 3rd instar larvae trachea, animals were dissected in cold PBS, and the isolated trachea were transferred to RNA Magic (BioBudget, Krefeld, Germany) and processed essentially as described earlier [19] with slight modifications. The tissue was homogenized using a Bead Ruptor 24 (BioLab products, Bebensee, Germany), and the RNA was extracted with the PureLink RNA Mini Kit (Thermo Fisher, Waltham, MA, USA) for phase separation with the RNA Magic reagent. An additional DNase treatment was performed following the on-column PureLink DNase treatment protocol (Thermo Fisher, Waltham, MA, USA). Sequencing libraries were constructed using the TruSeq stranded mRNA kit (Illumina, San Diego, USA) and 50 bp single-read sequencing was performed on an Illumina HiSeq 4000 with 16 samples per lane. Resulting sequencing reads were trimmed for low-quality bases and adapters using the fastq Illumina filter (http://cancan.cshl.edu/labmembers/gordon/fastq_illumina_filter/) and cutadapt (version 1.8.1)[61]. Transcriptomics analysis, including gene expression values and differential expression analysis, was done using CLC Genomics Workbench. The detailed protocols can be obtained from the CLC Web site (http://www.clcbio.com/products/clc-genomics-workbench). *Drosophila melanogaster* reference genome (Release 6)[62] was used for mapping in this research. RNAseq data are deposited in the GEO database (GSE212132). Mouse and human orthologs of the Drosophila DEGs were identified using the DIOPT program package[32].

The transcription factor binding site enrichment and the Gene Ontology enrichment analyses of the differentially expressed genes were carried out using Pscan (http://159.149.160.88/pscan/) and online GO enrichment analysis (http://geneontology.org/), respectively. We chose -450-50 bases of the annotated transcription start site of the genes as the transcription factor binding sites for enrichment analysis. The analysis used the TFBS matrices available in the JASPAR databases (version Jaspar 2018_NR). Data were visualized using the Circos software (http://circos.ca/software/).

## Statistics and reproducibility

We did not use statistical methods to predetermine sample sizes, but the sample sizes used in this study are similar to or higher than those used in previous studies [29, 63]. Specific approaches to randomly allocate samples to groups were not used, and the experiments were not performed in a blinded design. No data were excluded from the analysis. Prism (GraphPad version 7) was used for statistical analyses, and the corresponding tests used are listed in the figure legends.

## Acknowledgments

We thank Britta Laubenstein and Christiane Sandberg for excellent technical assistance, Maria Leptin and Norbert Perrimon for flies.

## Data availability

RNAseq data are deposited in the GEO database (GSE212132). Any additional information required to reanalyze the data reported in this paper is available from the lead contact upon request.

