## Supplemental figure and tables for "A Goldilocks principle of JAK/STAT signaling governs airway epithelial homeostasis and stress adaptation in *Drosophila*"

**by**

Xiao Niu, Christine Fink, Kimberley Kallsen, Leizhi Shi, Viktoria Mincheva, Sören Franzenburg, Ruben Prange, Jingjing He, Anita Bhandari, Iris Bruchhaus, Holger Heine, Judith Bossen, Thomas Roeder

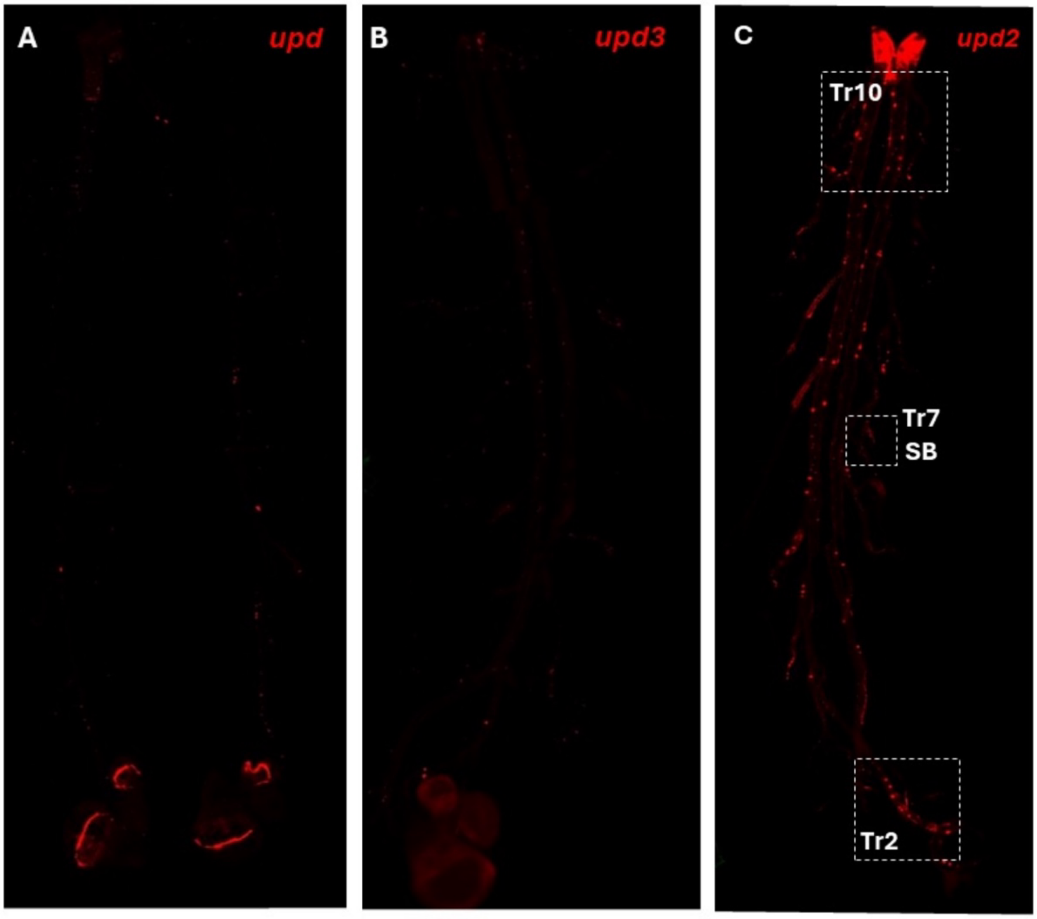


**Figure S1: JAK/STAT signaling activity in the larval trachea.** (A-C) Fluorescence micrographs of the trachea of ligand (*upd*, *upd2,* or *upd3*)-*Gal4>UAS-LacZ.nls* larvae stained to show ligand-expressing cells (anti-ß-galactosidase, red) and JAK/STAT signaling-activated cells (anti-GFP, green). *Upd2* displayed a higher transcript level in the trachea than *upd* (strong expression in other tissues, like imaginal discs) and *upd3*.

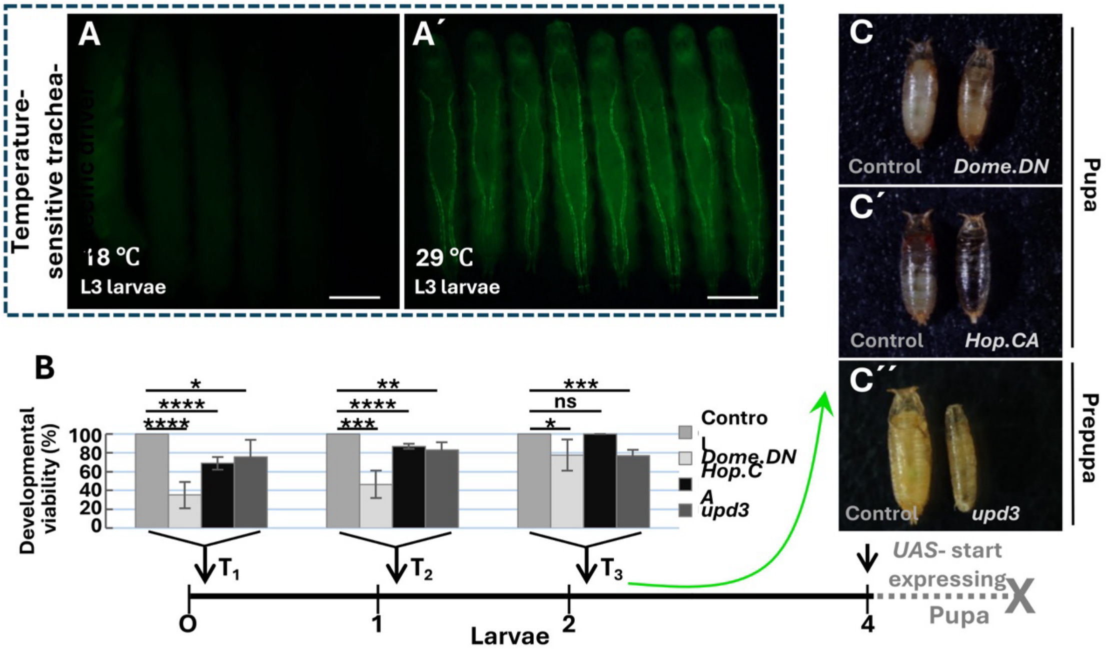


**Figure S2: Detection of heat-induced trachea-specific driver activity and the influence of JAK/STAT-specific components on developmental stages.** (A) The GFP signal reflects the effect in larvae expressing the heat-inducible driver *btl-Gal4, UAS-GFP; tub-Gal80[ts]* (*btl.ts*) reared at 18 ℃ (non-permissive, A) and 29 ℃ (permissive, A'). (B) Developmental viability of larvae with different genotypes (including *Dome.DN*, *Hop.CA,* and *upd3* expression in the trachea driven by *btl.ts*) and different starting points (T1-T3) of expression (indicated by black arrows) are shown. Statistical analysis of survival times is included, with four replicates performed, each containing 30 larvae in B. (C) None of these manipulations allowed for survival up to adulthood.

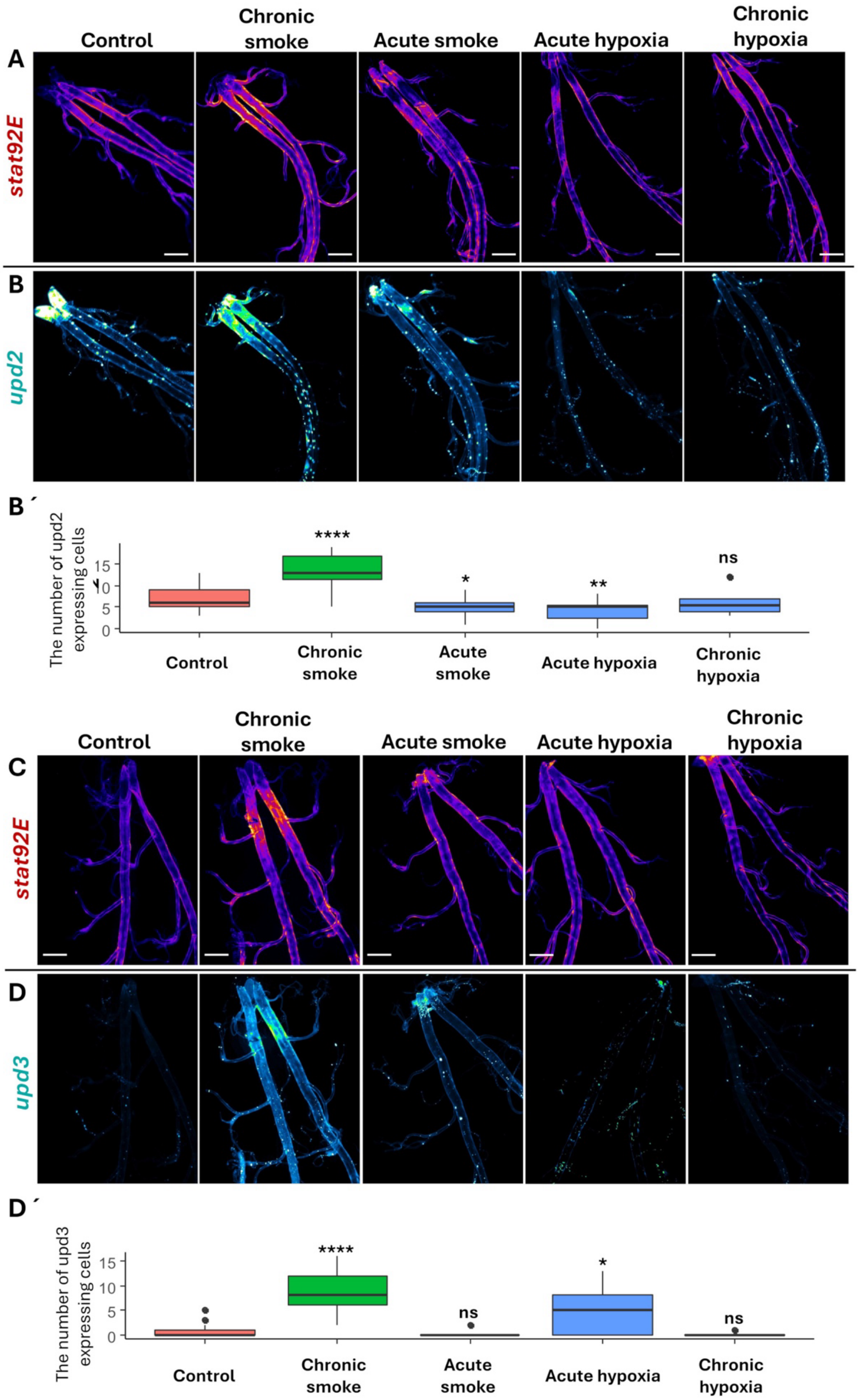
Control
Chronic smoke
Acute smoke
Acute hypoxia
Chronic hypoxia
A
stat92E
B
upd2
B´
The number of upd2 expressing cells
15
10
5
0
****
*
**
ns
Control
Chronic smoke
Acute smoke
Acute hypoxia
Chronic hypoxia
Control
Chronic smoke
Acute smoke
Acute hypoxia
Chronic hypoxia
C
stat92E
D
upd3
D´
The number of upd3 expressing cells
15
10
5
0
****
ns
*
ns
Control
Chronic smoke
Acute smoke
Acute hypoxia
Chronic hypoxia

**Figure S3: JAK/STAT signaling pathway activity under stress conditions.** Activity of the JAK/STAT pathway (*stat92E*; A, C; green) and the expression of its ligands, *upd2* (B) or *upd3* (D), in the trachea of larvae exposed to smoke and hypoxia. (A) Fluorescence micrographs of the trachea of larvae that were exposed for 2 days to smoke (chronic smoke), heavy smoke (acute smoke), strong hypoxia (acute hypoxia), and 2 days of hypoxia (chronic hypoxia). Trachea were stained for GFP (red) and Beta-galactosidase (green, cells that expressed *upd2/3*). (B', D') Number of cells expressing *upd2* (B') or *upd3* (D') under these different conditions. n=40, ns = not significant, * $p < 0.05$, ** $p < 0.01$; Student's t-test. Scale bar: 200 μm.

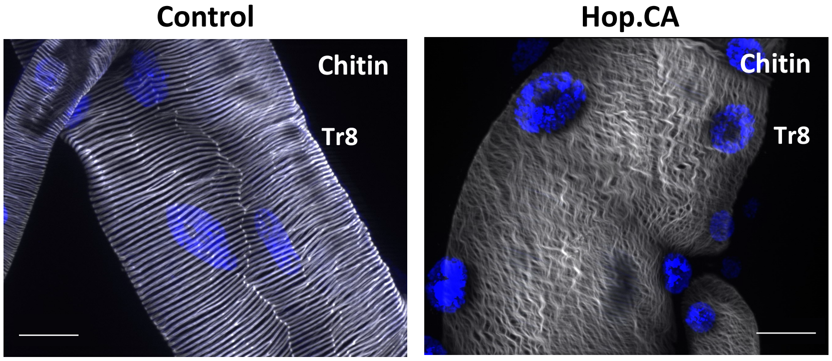


**Figure S4: Ectopic Hop.CA expression induced irregularities in the chitinous intima of the larval trachea.** Hop.CA driven by btl-Gal4 induced changes in the chitinous intima of the trachea, especially destroying the highly ordered orientation of taenidial rings. Chitin was stained with Calcofluor LRP. Blue, DAPI staining of nuclei. Scale bar: 20 μm.

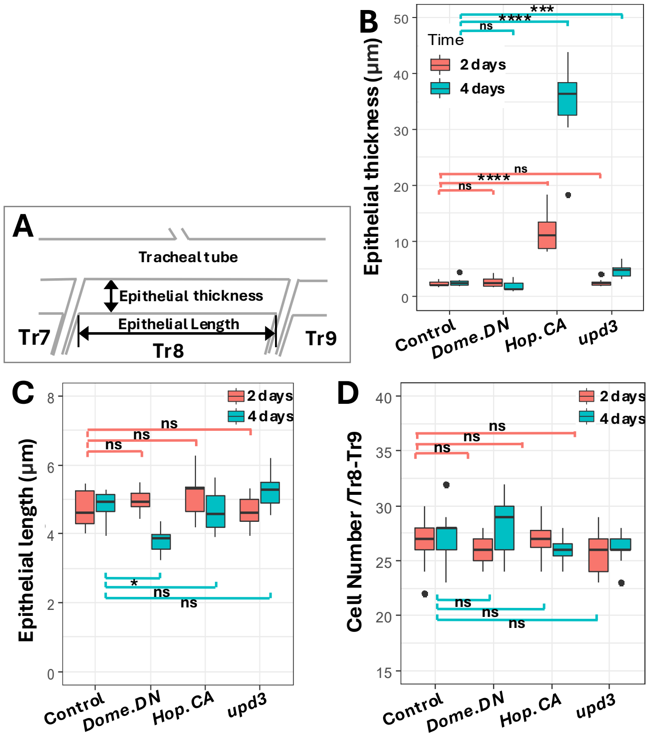

**Figure S5: Hop.CA activation induces only an epithelial thickening.** (A) Illustration of epithelial thickness and epithelial length measurement (arrows). (B-D) Quantification of tracheal epithelial thickness (B), epithelial length (C), and the cell number (D) of larvae with signaling pathway manipulation driven by *btl.ts-Gal4* for 2 or 4 days.

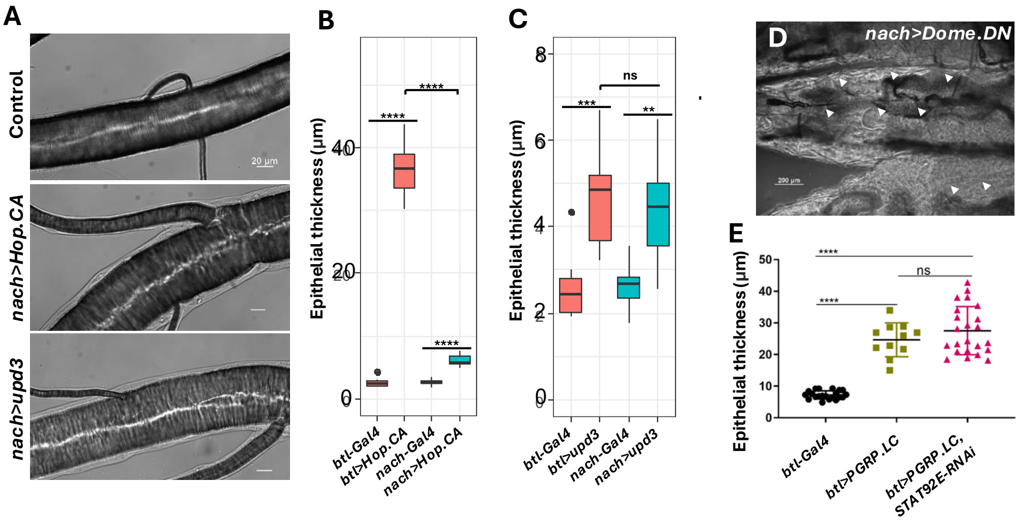


**Figure S6: Ectopic JAK/STAT activation with weaker driver lines induced weaker phenotypes.** (A-C) Mild activation (driven by *nach-Gal4*) of epithelial JAK/STAT (*Hop.CA* or *upd3*) signaling mitigated the thickening phenotype. (A) Micrographs of trachea from larvae with JAK/STAT activation. Scale bar: 20 µm. (B, C) Quantification of the epithelial thickness (DT8). (D) Micrographs of trachea from larvae experiencing suppression of the JAK/STAT pathway (*nach>Dome.DN*). Triangles indicate the tracheal position. Scale bar: 200 µm. (E) Quantification of the epithelial thickness (DT8) of *btl.ts-Gal4*, *btl.ts>PGRP.LC* and *btl.ts>PGRP.LC;STAT92E-RNAi* larvae. Induction of expression in the 3rd instar larval stage for one day. Green arrow: defective region, ns = not significant, * $p < 0.05$, ** $p < 0.01$, *** $p < 0.001$, **** $p < 0.0001$; Student's t-test.

Table S1: Kegg pathway analysis of genes, whose expression is regulated in response to *Hop.CA* overexpression.

| Term | # regulated gene | #REF | corrective p-value |
|---|---|---|---|
| Protein processing in endoplasmic reticulum | 37 | 122 | 0.000196415 |
| Metabolism of xenobiotics by cytochrome P450 | 21 | 62 | 0.005458041 |
| Glutathione metabolism | 20 | 62 | 0.006987233 |
| Drug metabolism - cytochrome P450 | 20 | 62 | 0.006987233 |
| Metabolic pathways | 143 | 924 | 0.010283682 |
| Fatty acid metabolism | 14 | 42 | 0.033601606 |
